# Mice with *GNAO1* R209H Movement Disorder Variant Display Hyperlocomotion Alleviated by Risperidone

**DOI:** 10.1101/662031

**Authors:** Casandra L. Larrivee, Huijie Feng, Josiah A. Quinn, Vincent S. Shaw, Jeffrey R. Leipprandt, Elena Y. Demireva, Huirong Xie, Richard R. Neubig

**Affiliations:** Department of Comparative Medicine and Integrative Biology, Michigan State University, 784 Wilson Rd, East Lansing MI, 48824 US; Department of Pharmacology & Toxicology, Michigan State University, 1355 Bogue Street, East Lansing, MI 48824 US; Transgenic and Genome Editing Facility, Institute for Quantitative Health Science and Engineering, Michigan State University, 775 Woodlot Drive, East Lansing MI 48823 US; Nicholas V. Perricone, M.D., Division of Dermatology, Department of Medicine, Michigan State University, 1355 Bogue Street, East Lansing, MI 48824

**Keywords:** *GNAO1*, Movement Disorder, Risperidone, Rare Disease

## Abstract

Neurodevelopmental disorder with involuntary movements (NEDIM, OMIM: 617493) is a severe, early onset neurological condition characterized by a delay in psychomotor development, hypotonia, and hyperkinetic involuntary movements. Heterozygous *de novo* mutations in the *GNAO1* gene cause NEDIM. Gα_o_, the gene product of *GNAO1,* is the alpha subunit of G_o_, a member of the heterotrimeric G_i/o_ family of G-proteins. G_o_ is found abundantly throughout the brain but the pathophysiological mechanisms linking Gα_o_ functions to clinical manifestations of *GNAO1-*related disorders are still poorly understood. One of the most common mutant alleles among the *GNAO1* encephalopathies is the c.626G>A or p.Arg209His (R209H) mutation. We developed heterozygous knock-in *Gnao1*^*+/*R209H^ mutant mice using CRISPR/Cas9 methodology to assess whether a mouse model could replicate aspects of the NEDIM clinical pattern. Mice carrying the R209H mutation exhibited increased locomotor activity and a modest gait abnormality at 8-12 weeks. In contrast to mice carrying other mutations in *Gnao1*, the *Gnao1*^*+/R209H*^ mice did not show enhanced seizure susceptibility. Levels of protein expression in multiple brain regions were unchanged from WT mice but the nucleotide exchange rate of mutant R209H Gα_o_ was 9 times faster than WT. The atypical neuroleptic risperidone has shown efficacy in a patient with the R209H mutation. It also alleviated the hyperlocomotion phenotype observed in our mouse model but suppressed locomotion in WT mice as well. In this study, we show that *Gnao1*^*+/R209H*^ mice mirror elements of the patient phenotype and respond to an approved pharmacological agent.

## Introduction

Gα_o_ is the alpha subunit of the heterotrimeric G protein G_o_. It is the most abundant heterotrimeric G protein in the central nervous system comprising 1% of mammalian brain membrane protein. Mutations in *GNAO1*, which encodes Gα_o_, have been linked to two distinct neurological conditions. In 2013, four children with early infantile epileptic encephalopathy were identified with mutations in *GNAO1* (Nakamura et al., 2013). Since then, a growing number of patients presenting with epilepsy and/or hyperkinetic movement disorders have been found to exhibit *de novo* mutations in *GNAO1* (Feng, Khalil, Neubig, & Sidiropoulos, 2018; Feng et al., 2017). It was recognized in 2016 that some *GNAO1* mutations result in movement disorders without epilepsy (Kulkarni, Tang, Bhardwaj, Bernes, & Grebe, 2016; Saitsu et al., 2016). To date there are over 70 published cases of children with mutations in *GNAO1* presenting with early infantile epileptic encephalopathy (EIEE17: OMIM 615473) and/or neurodevelopmental disorder with involuntary movements (NEDIM: OMIM 617493) (Ananth et al., 2016; Arya, Christine, L., L., & D., 2017; Blumkin et al., 2018; Bruun, 2018; E.-R. Consortium, Project, & Consortium, 2014; E. K. Consortium, 2016; Danti et al., 2017; Dhamija, W., B., & P., 2016; Gawlinski et al., 2016; Gerald et al., 2018; Honey et al., 2018; Koy et al., 2018; Kulkarni et al., 2016; Law et al., 2015; Marcé-Grau et al., 2016; Marecos, S., I., E., & A., 2018; Menke et al., 2016; Meredith et al., 2019; Nakamura et al., 2013; Okumura et al., 2018; Rim et al., 2018; Saitsu et al., 2016; Sakamoto et al., 2017; Sanem et al., 2016; Schirinzi et al., 2018; Schorling et al., 2017; Takezawa et al., 2018; Talvik et al., 2015; Waak et al., 2018; Xiong et al., 2018).

More than forty pathological variants of *GNAO1* have been reported. Using a cell-based biochemical signaling assay, we classified many of those Gα_o_ variants for their ability to support inhibition of cAMP production in transfected HEK293 cells (Feng et al., 2017). Some mutant Gα_o_ proteins were unable to support receptor-mediated inhibition of cAMP which classifies them as having a loss-of-function (LOF) mechanism *in vitro*. The LOF mutants were associated with epilepsy (Feng et al., 2017). In contrast, mutations resulting in enhanced cAMP inhibition (gain-of-function, GOF) or those which supported normal cAMP regulation (NF) were generally associated with movement disorders, though some patients with these mutations also have a relatively mild seizure disorder(Feng et al., 2017).

To permit mechanistic studies and preclinical drug testing, we had previously created a mouse model with a *GNAO1* GOF mutation, G203R, that was identified in patients who showed both epilepsy and movement disorders (Saitsu et al., 2016). As predicted, the *Gnao1*^+/G203R^ (G203R) mutant mice exhibited motor coordination and gait abnormalities as well as enhanced seizure susceptibility in pentylenetrazol (PTZ) kindling studies (Feng et al., 2019). The R209H mutations is one of the most common pathogenic *GNAO1* mutations (Schirinzi et al., 2018). Patients with *de novo*, heterozygous R209H mutations in *GNAO1* display severe choreoathetosis and dystonia but do not exhibit seizures (Ananth et al., 2016; Dhamija et al., 2016; Kulkarni et al., 2016; Marecos et al., 2018; Menke et al., 2016; Meredith et al., 2019). Interestingly, the R209H mutation was found to have essentially normal function for cAMP inhibition in HEK 293T cells. Despite this normal function in an *in vitro* assay, it causes a severe form of movement disorder in patients, often requiring intensive care unit admission (Ananth et al., 2016; Marecos et al., 2018). This discrepancy between the normal functionality *in vitro* and its clear clinical pathologic role made the R209H mutation of substantial interest for *in vivo* physiological studies to better understand its mechanism.

We used a battery of behavioral tests to measure motor skills in heterozygous *Gnao1*^*+/R209H*^ mutant mice as well as performing PTZ kindling studies to assess seizure susceptibility. As expected *Gnao1*^*+/R209H*^ mice did not show enhanced seizure susceptibility in PTZ kindling studies. Male and female *Gnao1*^*+/R209H*^ mice both displayed significant hyperactivity in an open field assessment. This finding was surprising as mice in our previous *GNAO1*-related movement disorder model, *Gnao1*^*+/G203R*^ did not show significant differences on the open field test (Feng et al., 2019). This difference in movement phenotype is consistent with the wide heterogeneity of movement patterns displayed by patients with GNAO1 mutations (Ananth et al., 2016; Dhamija et al., 2016; Feng et al., 2018; Kulkarni et al., 2016; Marcé-Grau et al., 2016; Menke et al., 2016; Nakamura et al., 2013).

Having a mouse model with a strong movement phenotype allowed us to begin allele-specific preclinical drug testing and will facilitate mechanistic studies of *GNAO1* mutants. The neuroleptic risperidone was reportedly beneficial in a patient a *GNAO1* R209H mutation (Ananth et al., 2016). Here we show that risperidone attenuates hyperactivity in our R209 Hmutant mice. This suggests that risperidone or related agents may be beneficial for *GNAO1* patients with the R209H mutation.

## Experimental Procedures

### Animals

*Gnao1*^*+/R209H*^ mice on a C57BL/6J background were generated in the MSU Transgenic and Gene Editing Facility (https://tgef.vprgs.msu.edu) as described below. Mice (8-12 weeks old) were housed on a 12-hour light/dark cycle, with *ad libitum* access to food and water. All experiments were performed in accordance with NIH guidelines and protocols approved by the Michigan State University Institutional Animal Care and Use Committee.

### Generation of Gnao1 R209H edited mice

Mutant *Gnao1*^*+/R209H*^ mice were generated via CRISPR/Cas9 genome editing on a C57BL/6J genomic background. CRISPR gRNA selection and locus analysis were performed using the Benchling platform (Benchling, Inc. San Francisco, CA.). A gRNA targeting exon 6 of the *Gnao1* locus (ENSMUSG00000031748) was chosen to cause a double strand break (DSB) 3bp downstream of codon R209. A single stranded oligodeoxynucleotide (ssODN) carrying the R209H mutation CGC > CAC with short homology arms was used as a repair template (Figure 1 and Table 1). Ribonucleoprotein (RNP) complexes consisting of a synthetic crRNA/tracrRNA hybrid and Alt-R^®^ S.p. Cas9 Nuclease V3 protein (Integrated DNA Technologies, Inc. Coralville, IA), were used to deliver CRISPR components along with the ssODN to mouse zygotes via electroporation as previously described (Feng et al., 2019; Feng et al., 2017). Edited embryos were implanted into pseudo-pregnant dams using standard techniques. Resulting litters were screened by PCR (Phire Green HSII PCR Mastermix, F126L, Thermo Fisher, Waltham, MA.), T7 Endonuclease I assay (M0302, New England Biolabs Inc.) and Sanger sequencing (GENEWIZ, Inc. Plainfield, NJ) for edits of the target site.

**Table 1.**
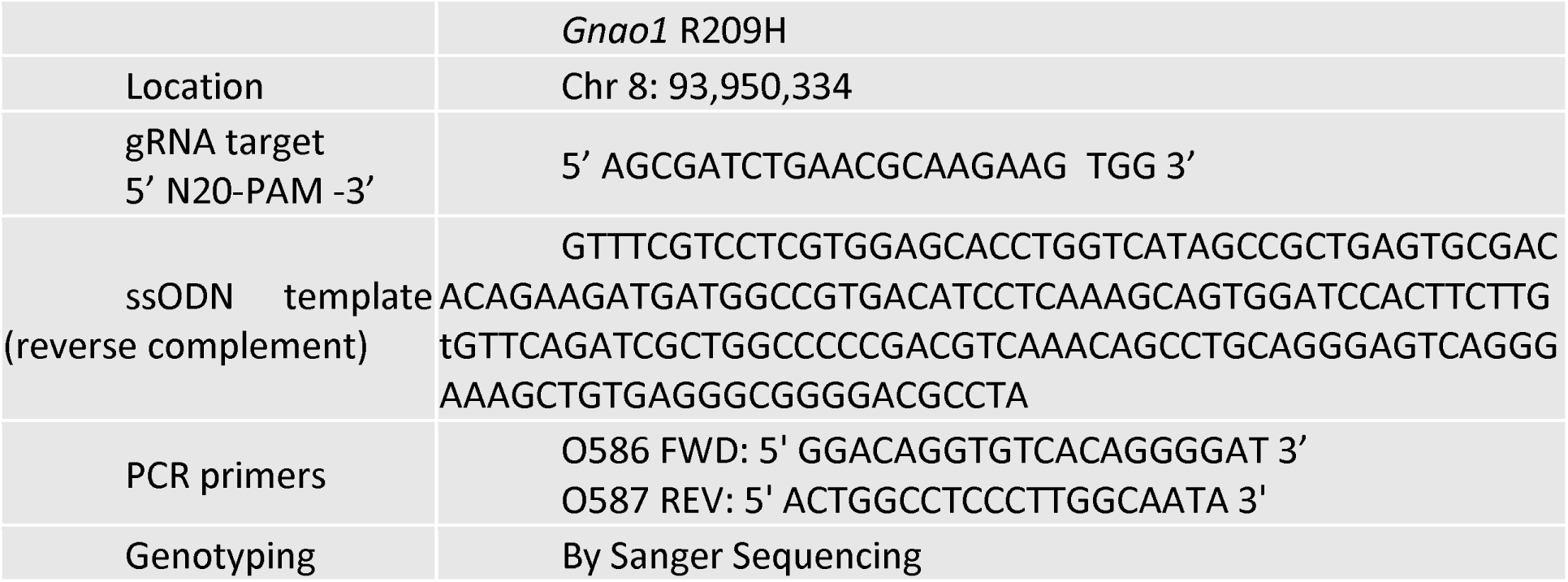
Location and sequence of gRNA and ssODN template for CRISPR-Cas targeting Gnao1 locus; primers and genotyping method for *Gnao1*^*+/R209H*^ mice.

**Figure 1.**
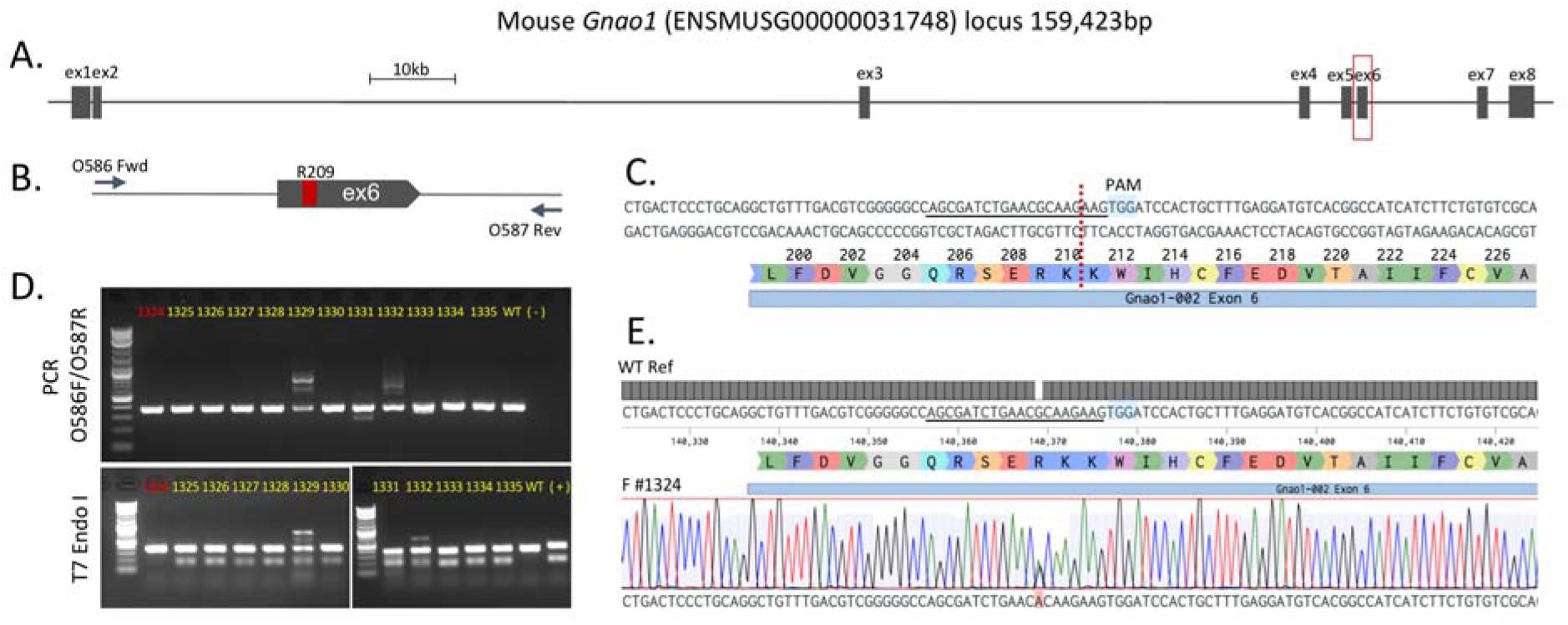
Targeting of the mouse *Gnao1* locus. (A) Mouse *Gnao1* genomic locus (exon size not to scale), red outline is magnified in (B) showing exon 6 and relative location of codon 209 and PCR primers O586 and O587. (C) Location and exact sequence of gRNA target within exon 6, dotted red line denotes DSB, PAM is highlighted and sequence corresponding to gRNA protospacer is underlined (also in E). (D) Raw gel electrophoresis images showing PCR of the target region and T7 Endonuclease I (T7 Endo I) digestion analysis of founders 1324 – 1335 (n=12), with WT, H2O (-) and T7Endo I (+) controls. Founder 1324 (red number) was positive for the mutation on one allele and WT on the other, note that the single bp mismatch was not reliably detected by T7 Endo I assay. (E) Exact sequence of edited founder 1324 as aligned to WT reference genome, two peaks (G and A) are detected on the sequence chromatogram, indicating the presence of both WT and edited R209H allele.

### Genotyping and Breeding

Studies were done on N1 R209H heterozygotes with comparisons to littermate controls. To generate *Gnao1*^*+/R209H*^ heterozygotes (N1 backcross), 2 founder *Gnao1*^*+/R209H*^ mice, 1 male and 1 female, were crossed with WT C57BL/6J mice obtained directly from The Jackson Laboratory (Bar Harbor, ME).

DNA was extracted by an alkaline method (26) from ear clips done before weaning. PCR products were generated with primers flanking the mutation site (Fwd 5’ GGACAGGTGTCACAGGGGAT 3’; 5’ ACTGGCCTCCCTTGGCAATA 3’) which produces a 375 base pair (bp) product. Reaction conditions were: 0.8 μl template, 4 μl 5x Promega PCR buffer, 0.4 μl 10mM dNTPs, 1 μl 10 μ M Forward Primer, 1 μl 10 μ M Reverse Primer, 0.2 μl Promega GoTaq and 12.6 μ l DNase free water (Promega catalog # M3005, Madison WI). Samples were denatured for 4 minutes at 95° C then underwent 32 cycles of PCR (95° C for 30 seconds, 63°C for 30 seconds, and 72°C for 30 seconds) followed by a 7 minute final extension at 72°C. Ethanol precipitation was done on the PCR products and then samples were sent for Sanger sequencing (GENEWIZ, Inc. Plainfield, NJ).

### Behavioral Assessment

Male and female *Gnao1*^*+/R209H*^ mice (8-12 weeks of age) and their *Gnao1*^*+/+*^ littermates underwent a battery of behavioral testing to assess motor phenotype as described previously (Feng et al., 2019). Before each experiment, mice were acclimated for 10 minutes to the testing room. Experiments were performed by two female researchers. All behavioral studies were done by individuals who were blind to the genotype of the animals until completion of data collection.

### Open Field

The open field test is frequently used to assess locomotion, exploration and anxiety (Feng et al., 2019; Seibenhener & Wooten, 2015; Tatem et al., 2014). The test was conducted in Fusion VersaMax clear 42 cm × 42cm × 30cm arenas (Omnitech Electronics, Inc., Columbus, Ohio). *Gnao1*^*+/R209H*^ mice of both sexes and their littermates were placed in the arena for 30 minutes. Using the Fusion Software, we evaluated distance traveled (cm) in terms of novelty, sustained, and total movement corresponding to the first 10 minutes, 10-20 minutes, and total of 30 minutes. As a potential measure of anxiety, the fraction of time spent in the center was assessed. The center area was defined as the 20.32cm × 20.32cm area within the middle of the arena.

### Rotarod

To assess motor skills, we used the Economex accelerating RotaRod (Columbus Instruments, Columbus OH). The protocol occurred over a two-day period. On day 1, mice were trained for three 2-min training sessions, with 10 minutes between each training trial. During the first two sessions, the rotarod maintained a constant rotational speed of 5 rpm. The third training trial started at 5 rpm and accelerated at 0.1 rpm/sec for 2 minutes. The following day, mice ran three more trials with a 10-min break in between: two more 2-min training trials and a final 5-min test trial. Each of these trials started at 5 rpm with constant acceleration of 0.1 rpm/sec. For all training and test trials, latency to fall off the spindle was recorded in seconds.

### Grip Strength

To assess mouse grip strength, we used seven home-made weights (10, 18, 26, 34, 42, 49, 57 grams) with a 2.54 cm ring for the mouse to grasp. The mouse was held by the middle/base of the tail and lowered to the weight. Once the mouse grasped the weighted ring with its forepaws, the mouse was lifted until the weight cleared the bench. For each weight, the mouse was given up to three trials to suspend the weight above the table for 3 seconds. If cleared, the next heaviest weight was tried. If the weight wasn’t held, the total time and maximum weight lifted was recorded and a grip strength score calculated from (Deacon et al., 2016). The calculated score was normalized to mouse body weight which was measured the day of the test.

### DigiGait

Mouse gait analysis was performed on a DigiGait apparatus (Mouse Specifics, Inc, Framingham, MA). After acclimation, each mouse was placed on the treadmill at speeds of 18, 20, 22, 25, 28, 32, 36 cm/s. A 10-second clip was recorded with a video camera located below the belt. There was a 5-min rest between each speed. Recordings were analyzed with the DigiGait analysis program to assess the pre-specified parameters of stride length and paw angle variability. Values for all four paws were averaged to give one value per mouse – the reported n values are the number of mice. In addition, the maximum speed at which each mouse was able to successfully complete a 10-second test after three attempts was recorded as described (Feng et al., 2019).

### PTZ Kindling Study

A PTZ kindling protocol was performed as described (Feng et al., 2019; Kehrl et al., 2014) to assess mouse susceptibility to seizure induction. Mice were injected with a sub-convulsive dose of PTZ (40 mg/kg, i.p.) every other day for up to 24 days mice then were observed for 30 minutes post-dose. Kindling was defined as death or tonic-clonic seizures on two consecutive injection days after which mice were euthanized. Kaplan-Meier survival analysis was done based on the number of injections to achieve kindling.

### Gα_o_ *protein expression*

Mice (6-8 weeks old) were sacrificed and their brains were dissected into different regions and flash-frozen in liquid nitrogen. For Western Blot analysis, tissues were thawed on ice and homogenized for 5 min with 0.5 mm zirconium beads in a Bullet Blender (Next Advance; Troy, NY) in RIPA buffer (20mM Tris-HCl, pH7.4, 150mM NaCl, 1mM EDTA, 1mM β-glycerophospate, 1% Triton X-100 and 0.1% SDS) with protease inhibitor (Roche/1 tablet in 10 mL RIPA). Sample homogenates were centrifuged for 5 min at 4°C at⍰13,000 G. Supernatants were collected and protein concentrations determined using the bicinchoninic acid method (BCA method; Pierce; Rockford, IL). Protein concentration was normalized for all tissues with RIPA buffer and 2x SDS sample buffer containing β-mercaptoethanol (Sigma-Aldrich) was added. Thirty µg of protein was loaded onto a 12% Bis-Tris SDS-PAGE gel and samples were separated for 1.5 hrs at 160V. Proteins were then transferred to an Immobilon-FL PVDF membrane (Millipore, Billerica, MA) on ice either for 2 h at 100 V, 400 mA or overnight at 30V, 50mA. Immediately after transfer, PDVF membranes were washed and blocked in Odyssey PBS blocking buffer (Li-Cor) for 40 min at RT. The membranes were then incubated with anti-Gαo (rabbit; 1:1,000; sc-387; Santa Cruz biotechnologies, Santa Cruz, CA) and anti-actin (goat; 1:1,000; sc-1615; Santa Cruz) antibodies diluted in Odyssey blocking buffer with 0.1% Tween-20 overnight at 4°C. Following four 5-min washes in phosphate-buffered saline with 0.1 % Tween-20 (PBS-T), the membrane was incubated for 1 hr at room temperature with secondary antibodies (both 1:10,000; IRDye^®^ 800CW Donkey anti-rabbit; IRDye^®^ 680RD Donkey anti-goat; LI-COR Biosciences) diluted in Odyssey blocking buffer with 0.1 % Tween-20. The membrane was subjected to four 5-min washes in PBS-T and a final rinse in PBS for 5 min. The membrane was kept in the dark and the infrared signals at 680 and 800nm were detected with an Odyssey Fc image system (LI-COR Biosciences). The Gα_o_ polyclonal antibody recognizes an epitope located between positions 90-140 Gα_o_ (Santa Cruz, personal communication) so there should be no interference from the R209H mutation.

### Kinetics of nucleotide binding

To estimate GDP release rates, which can control activation of G proteins, we measured the kinetics of binding of the fluorescent ligand BODIPY FL GTPγS as described previously (McEwen, Gee, Kang, & Neubig, 2002). In brief, WT and mutant his6-tagged Gα_o_ subunits were expressed in E coli and purified by Ni-NTA affinity chromatography as previously described (Lee, Linder, & Gilman, 1994) and stored frozen at −80 C in 50 mM HEPES 100 mM NaCl buffer containing 50 uM GDP to stabilize the protein. Protein was diluted to 200 nM in binding buffer (50 mM HEPES, 10 mM MgCl2, 1 mM EDTA and 1 mM DTT, pH 8.0) then 100 nM BODIPY-GTPγS was added and fluorescence monitored at 485 nm ex, 510 nm em in a Tecan Infinite M1000 Pro microplate reader at 24 °C. Experiments were done on three different days each in duplicate and results averaged. Rate constants and t_1/2_ were determined by fitting results to the equation Y = Y0 + (Ymax-Y0)*(1-e^-k*t^) in GraphPad Prism v. 8.

### Risperidone effects on motor behavior

Naïve 8-12 week old *Gnao1* ^*+/R209H*^ and *Gnao1* ^*+/+*^ littermates of either sex were tested for effects of risperidone on their activity in the open field arena. The study was run over 5 days: On day 1, mice underwent the open field protocol described above to establish a baseline. On day 3, mice were habituated in the experimental room for 10 min then given a single i.p. dose of 2.0 mg/kg risperidone (Cayman Chemical, Ann Arbor, MI) or vehicle control. Risperidone was dissolved in DMSO at a concentration of 5 mg/ml, Further dilutions were done in DI water. Thirty minutes following injection, mice were placed in the open field arena for a 30-minute testing time. On day 5, mice were retested in the same open field protocol without injection to assess drug washout. A week later the same protocol was followed to test the effects an i.p. dose of 0.5 mg/kg risperidone had on the WT and R209H mice.

### Statistical Analysis

Data were analyzed with unpaired Student’s *t*-test or Mantel-Cox, *two-way* ANOVA with Bonferroni corrections as appropriate using Graphpad Prism 7.0 (GraphPad; La Jolla, CA). A p < 0.05 was considered the cut off for significance throughout. Detailed discussion of statistical analyses can be found within figure legends.

## Results

### Gnao1^+/R209H^ mice are produced at the expected frequency and have normal viability

Two founder *Gnao1*^*+/R209H*^ mice, 1 male and 1 female, were crossed with C57BL/6J mice. Out of 98 offspring of a cross of *Gnao1*^*+/R209H*^ with WT mice, 51 heterozygotes and 47 WT were observed. *Gnao1*^*+/R209H*^ mice exhibit no overt postural or movement abnormalities or seizures under normal housing conditions. Adult mice showed no statistically significant differences in weight between WT and *Gnao1*^*+/R209H*^ genotypes of either sex.

### Both male and female Gnao1^+/R209H^ show significant hyperactivity in the open field arena but no significant differences on the rotarod or grip strength test

Patients with R209H mutations present with hyperkinetic movement disorders (Ananth et al., 2016; Dhamija et al., 2016; Menke et al., 2016). To detect motor abnormalities, *Gnao1*^*+/R209H*^ mice were subjected to a battery of behavioral tests. The open field arena was used to test overall locomotor activity. We divided the test into two sections reflecting activity in a novel environment (0-10 minutes) then sustained activity (10-30 min). *Gnao1*^*+/R209H*^ mice of both sexes showed markedly increased activity in the sustained activity period compared to their wildtype littermates (Figure 2B). Females also had significantly increased activity in the first 10 minutes (novelty period, Fig. 2B). As a potential indicator of anxiety-like behavior. Male and female *Gnao1*^*+/R209H*^ mice also displayed reduced time in center (Figure 2A and 2B).

**Figure 2.**
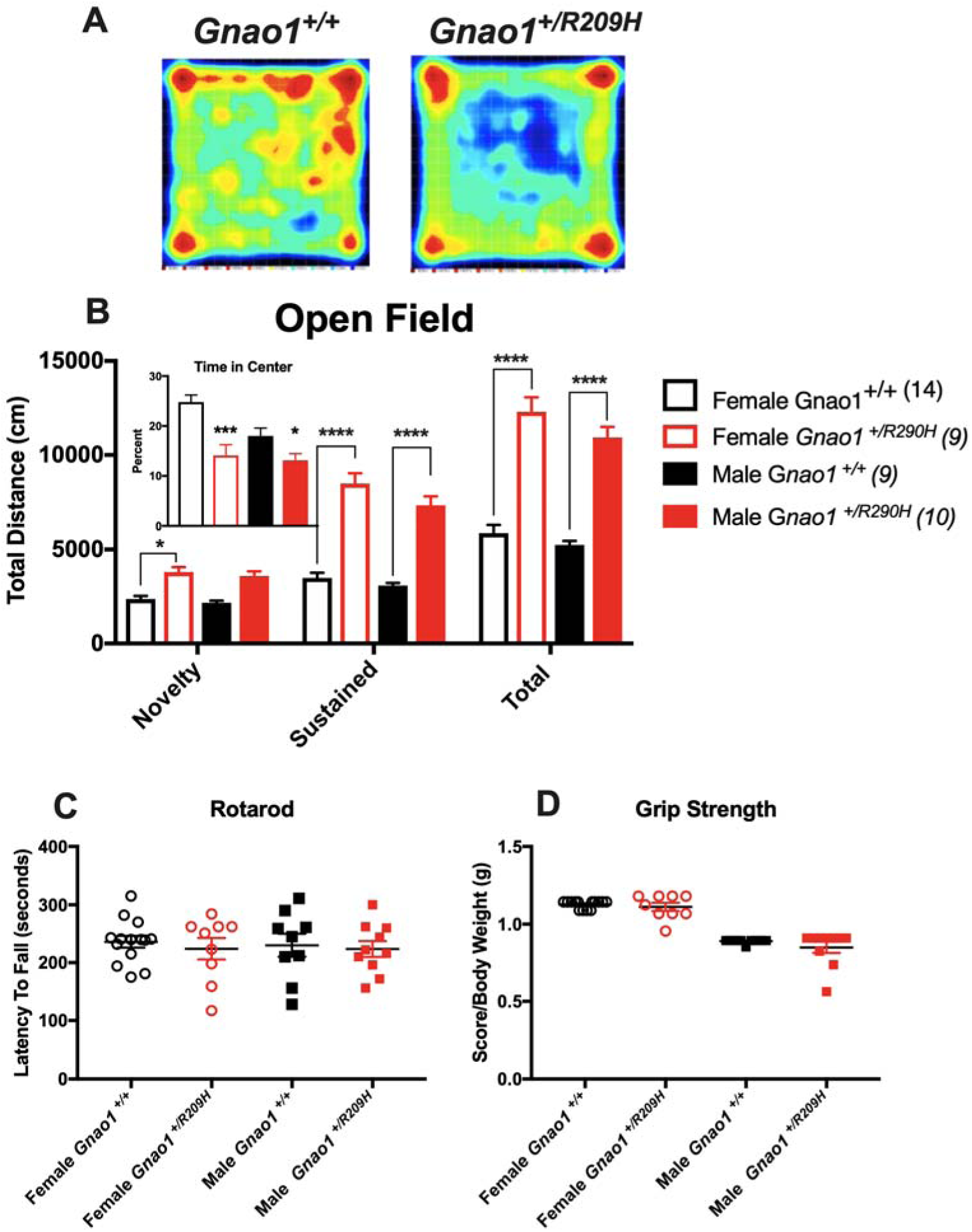
*Gnao1*^*+/R209H*^ mice show significant hyperactivity and reduced time in center in the open field arena. (A) Representative heat maps of *Gnao1*^*+/R209H*^ mice and *Gnao1*^*+/+*^ mice in the open field arena (B) Time spent in the open field arena was separated into 0-10 minutes (novelty) and 10-30 minutes (sustained). *Gnao1*^*+/R209H*^ male and female mice exhibit increased locomotion in the novelty period. Hyperactivity observed in the sustained period and for total travel time (2-way ANOVA; ****p < 0.0001, ***p < 0.001, * p < 0.05). Female R209H mutant mice also showed significantly greater distance traveled in the novelty period (*, P<0.05). *Gnao1*^*+/R209H*^ mice of both sexes spend less time in center areas of the open field arena compared to wildtype littermates. (C) Neither male nor female *Gnao1*^*+/R209H*^ mice show significant differences on the rotarod. (D) There is also no significant difference in grip strength between wildtype and *Gnao1*^*+/R209H*^ mice. Data are shown as mean ± SEM.

An accelerating rotarod test was used to assess motor coordination and balance. Neither male nor female *Gnao1*^*+/R209H*^ *mice* display impaired performance (Figure 2C). Also grip strength, showed no differences between *Gnao1*^*+/R209H*^ and wildtype littermates of either sex (Figure 2D).

### Male Gnao1^+/R209H^ mice display a modestly reduced stride length

Gait patterns were assessed using DigiGait analysis. Male *Gnao1*^*+/R209H*^ mice showed a highly significant genotype effect with reduced stride length compared to wildtype littermates (P<0.001, 2-way ANOVA) but the magnitude of the effect was modest (5.6% decrease averaged across all speeds). In post-test analysis, only the top speed (36 cm/sec) reached significance individually (p<0.01). Female *Gnao1*^*+/R209H*^ mice did not show significant differences in stride length from WT (Figure 3C & 3D). However as previously reported for G203R mutant mice, the female *Gnao1*^*+/R209H*^ showed a significantly reduced maximum run speed on the treadmill (Figure 3E p<0.01, t-test). This was not due to reduced body size as length, as detected by the digigait system, (WT 9.54 cm: vs R209H 10.17 cm), and weights were not significantly different. There was no significant difference in paw angle variability for either males or females.

**Figure 3.**
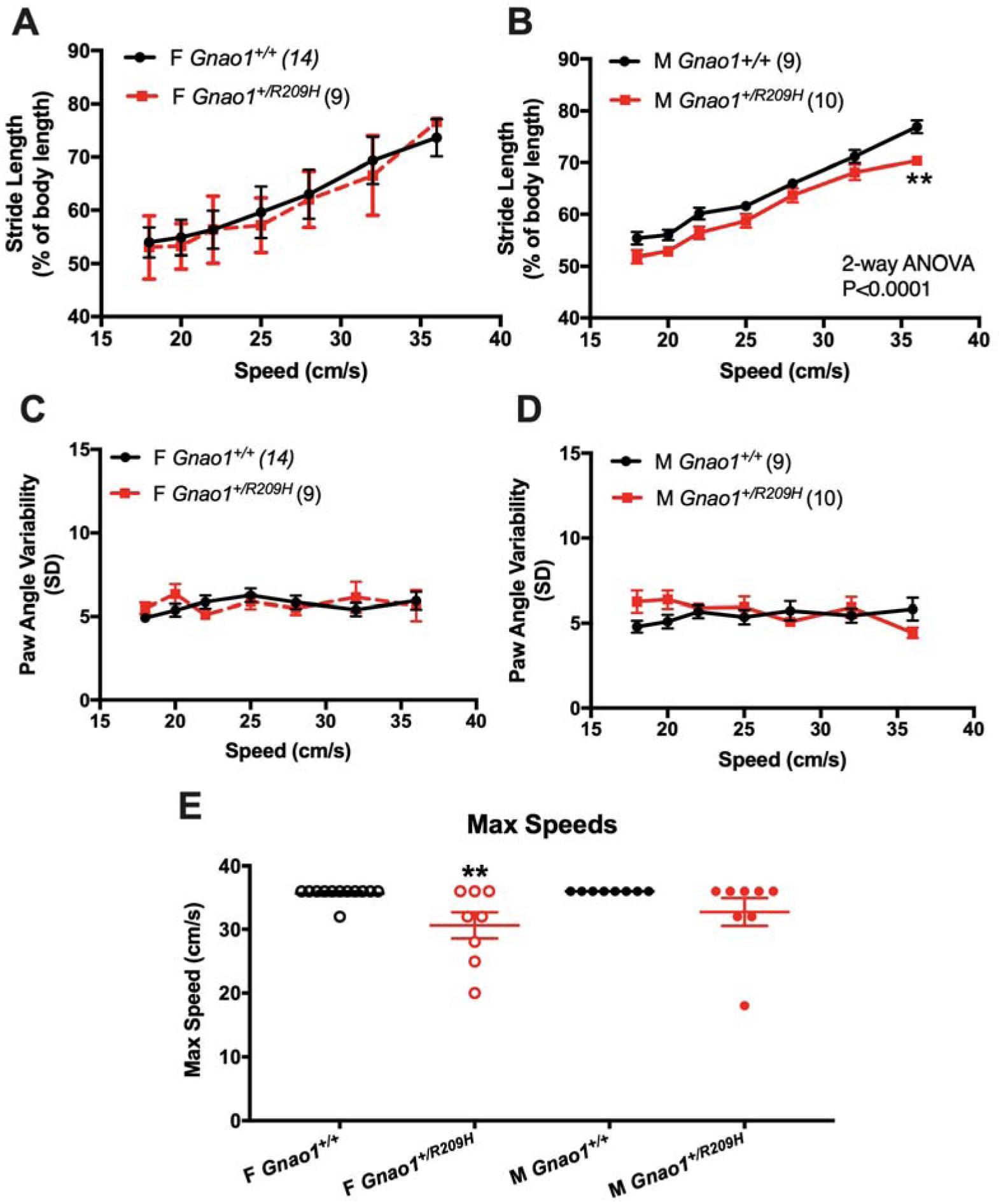
Male and female *Gnao1*^*+/R209H*^ mice shows gait abnormalities in different tests on the DigiGait imaging system. (A & B) Male *Gnao1*^*+/R209H*^ mice showed reduced stride length compared to wildtype littermates (2-way ANOVA with Bonferroni multiple comparison post-test), while female *Gnao1*^*+/R209H*^ mice showed a normal stride length. (C & D) Neither male nor female *Gnao1*^*+/R209H*^ exhibited significant differences in paw angle variability compared to wildtype littermates. (E) At speeds greater than 25 cm/s female *Gnao1*^*+/R209H*^ had a reduced ability to run on the treadmill.

### Gnao1^+/R209H^ mice are not sensitive to PTZ kindling

Repeated application of a sub-threshold convulsive stimulus leads to the generation of full-blown convulsions in a process called kindling (Dhir, 2012). *GNAO1* variants differ in their ability to cause epileptic seizures in patients. Those carrying the R209H mutant allele do not exhibit a seizure disorder (Ananth et al., 2016; Kulkarni et al., 2016; Menke et al., 2016). In accordance with the patients’ pattern, *Gnao1*^*+/R209H*^ mice did not show increased susceptibility to kindling-induced seizures (Figure 4A & 4B). This contrasts with our previous report of increased kindling sensitivity in male G203R and female G184S mutant mice (Dhamija et al., 2016; Feng et al., 2019; Kehrl et al., 2014).

**Figure 4.**
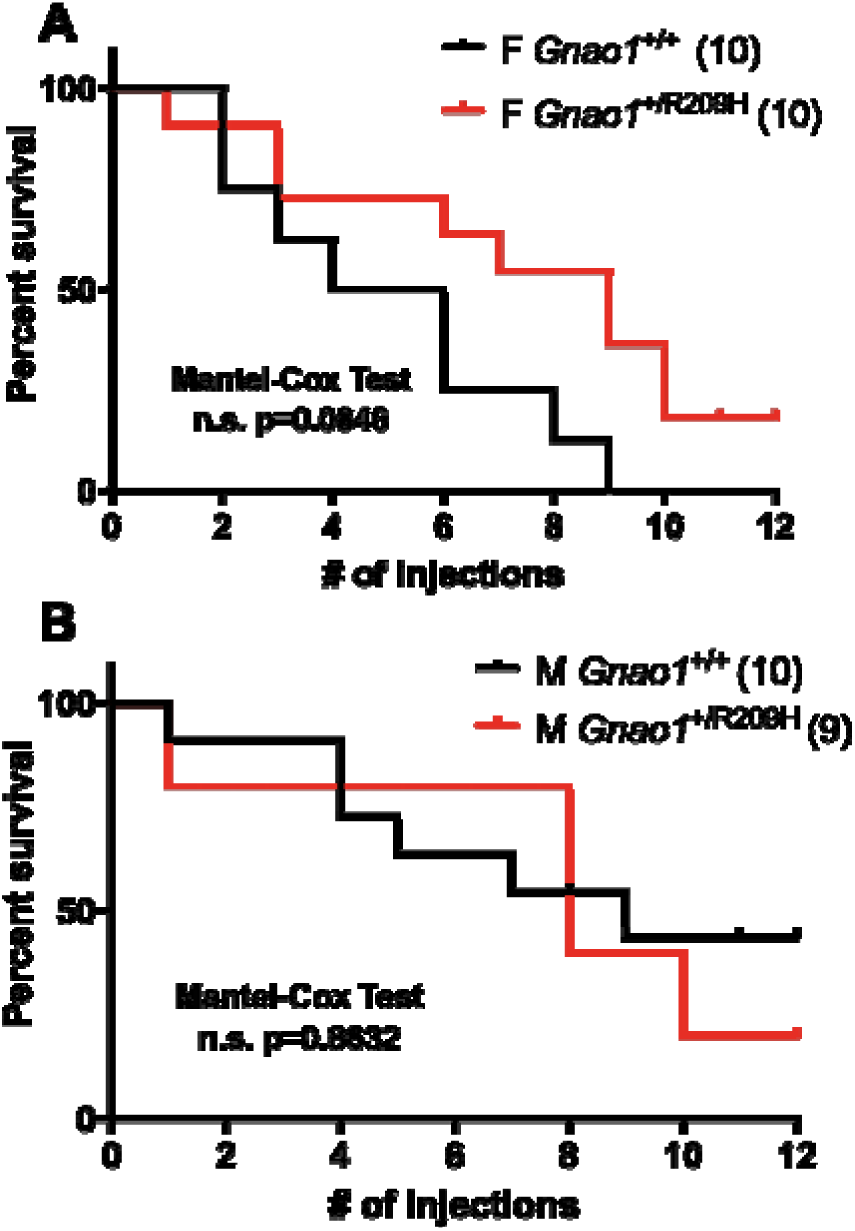
*Gnao1*^*+/R209H*^ mice do not have an enhanced pentylenetetrazol (PTZ) kindling response. (A&B) Neither male nor female *Gnao1*^*+/R209H*^ mice showed significant differences in sensitivity to PTZ injection compared to wildtype littermates (n.s. Mantel-Cox test).

### Gnao1^+/R209H^ mice have normal Gα_o_ protein expression in the brain

To understand why R209H mutant mice do not show a phenotypic difference in the kindling test while G203R mutants do, we assessed Gα_o_ protein expression levels. Cortex, hippocampus, striatum, cerebellum, brain stem and olfactory bulb were harvested and homogenized to measure the effect of the R209H mutation on Gα_o_ protein expression. Western blots showed no difference in Gα_o_ protein expression between WT and *Gnao1*^*+/R209H*^ mice in any of the measured brain regions (Figure 5A & 5B). This is consistent with our previous analysis of protein expression in HEK293T cells with transiently transfected Gα_o_ R209H plasmid (Feng et al., 2017). The normal expression of the R209H mutant contrasts with the reduced expression of Gα_o_ in brains of Gα_o_ ^+/G203R^ mutant mice (Feng et al., 2019).

**Figure 5.**
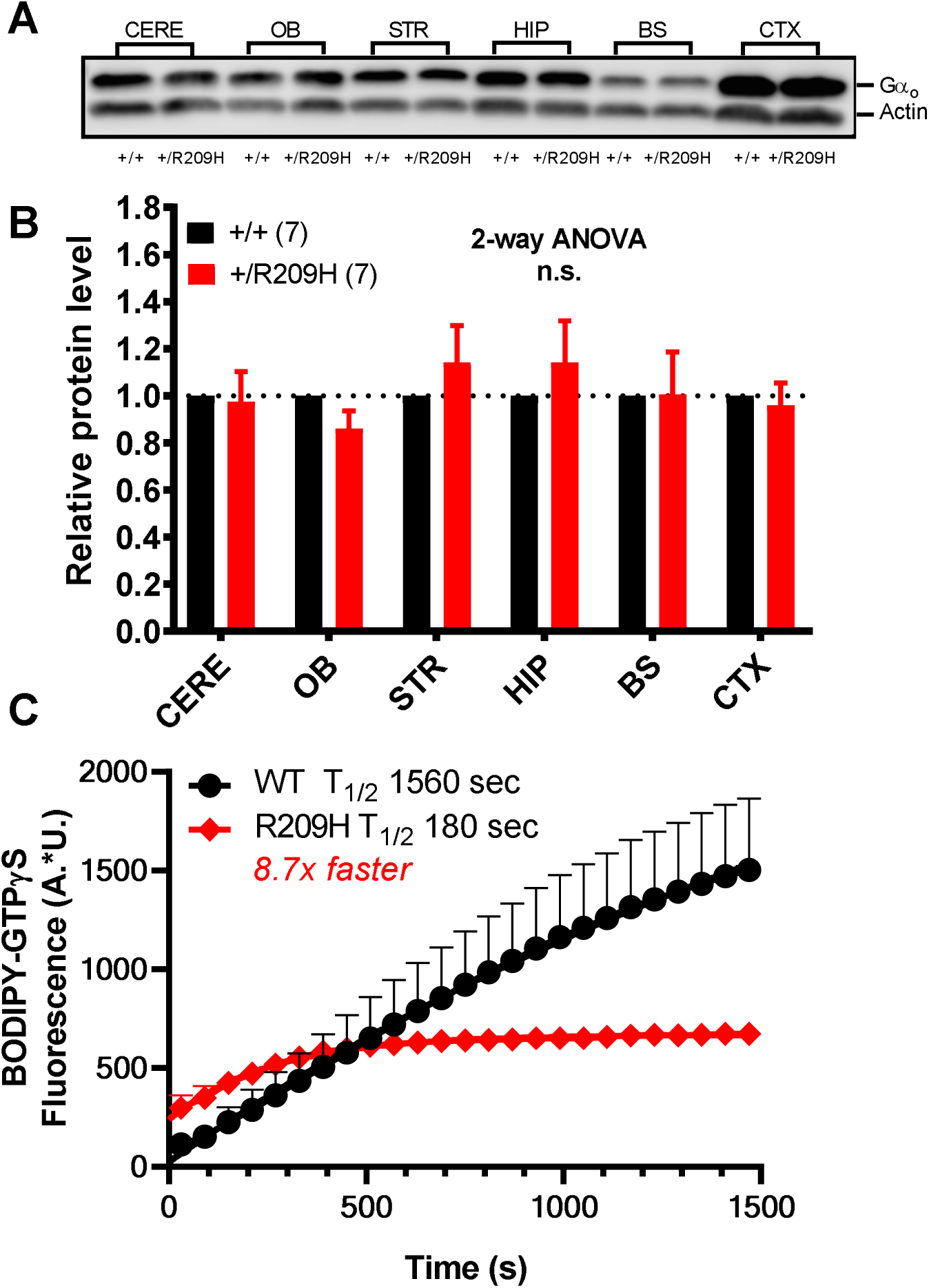
Biochemical analysis of brain Gα_o_ protein levels and in vitro nucleotide exchange for WT and R209H mutant G*α*_o_. A) Brain regions (cortex, hippocampus, striatum, cerebellum, brain stem and olfactory bulb homogenates) from WT and G*nao1*^*+/R209H*^ mice were assessed by Western blot for levels of Gαo protein. B) Protein levels of *Gnao1*^*+/R209H*^ brain samples were quantified using Li-Cor IRDye staining and values were normalized to actin levels and then to the control sample from WT run on the same day. There was no significant difference in any of the regions between WT and mutant mice (n=7). C) The kinetics of BODIPY-FL *GTP*γS binding were measured as described in Materials and Methods. The time course of the fluorescence increase was fit to an exponential function (connecting lines) and the T1/2 calculated from the rate constant of the exponential function (see methods). Nucleotide binding experiments were performed in duplicate and results from the separate days were averaged. Error bars and mean +-SEM (n=3). For some data points, error bars are smaller than the symbol and are not shown.

### Kinetics of nucleotide binding to mutant Gα_o_

Both patients and mice heterozygous for the Gα_o_ R209H mutation display movement phenotypes however, the mutant Gα_o_ supports normal cAMP regulation in the cell-based cAMP assay(Feng et al., 2017). To consider possible mechanisms for this discrepancy we measured the rate of GDP release from mutant Gα_o_ using the BODIPY-GTPγS binding kinetics method (McEwen et al., 2002). The rate of GTPγS binding to WT Gα_o_ was slow (T_1/2_ 1560 seconds, Fig. 5C) as expected. The R209H mutant Gα_o_ showed markedly faster binding of BODIPY-GTPγS (T_1/2_ 180 seconds 8.7x faster than WT) suggesting a faster rate of GDP release. In addition to the faster rate of binding, the amplitude of binding was lower and the initial fluorescence immediately after adding the GTPγS was slightly higher for the R209H mutant. While this lower amplitude of binding might be due to either a lower affinity of the mutant for the BODIPY-GTPγS or to instability under the binding conditions we do show that the R209H mutant definitely does have abnormal interactions with nucleotides – GDP and/or GTP.

### Risperidone treatment attenuated the hyperactivity of Gnao1^+/R209H^ mice

Patients with *GNAO1* mutations were tried on multiple drugs to alleviate motor symptoms (Table S1). Risperidone, an atypical antipsychotic drug showed beneficial effects in one of the patients. It has also been effective in drug-induced dyskinesia (Carvalho, Silva, Abílio, Barbosa, & Frussa-Filho, 2003). We show that *Gnao1*^*+/R209H*^ mice exhibit complete abrogation of movement at 2 mg/kg risperidone, which recovered upon retesting 2 days later (Figure 6A&C). WT mice also show a significant decrease in locomotion after 2 mg/kg risperidone treatment (Figure 6A). After a single 0.5 mg/kg dose of risperidone both WT and *Gnao1*^*+/R209H*^ mice exhibit a decrease in locomotion compared to vehicle treated littermates (Figure 6B). As expected, hyperactivity of mutant mice was observed during baseline testing on day 1 and the hyperactivity had returned on day 5 following the 2-day washout period after the risperidone doses (Figure 7C). Neither 2.0 mg/kg nor 0.5 mg/kg selectively affected *Gnao1*^*+/R209H*^ as assessed by percent suppression of distance travelled (Figure S1).

**Figure 6.**
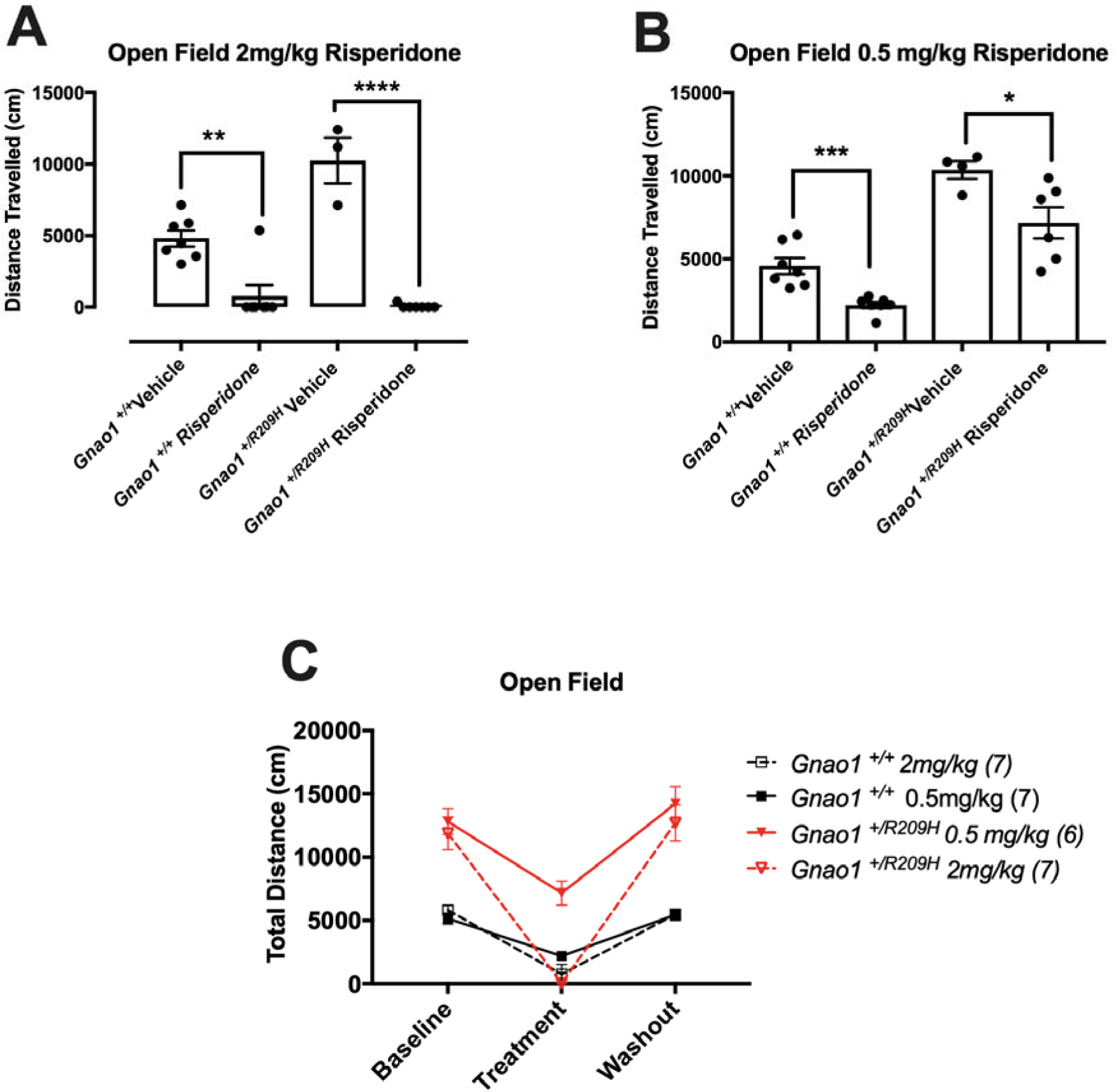
Risperidone treatments decreases hyperkinetic movements in *Gnao1* ^*+/R209*^. (A) *Gnao1*^*+/R209H*^ mice show complete abrogation of movement compared to vehicle treated *Gnao1*^*+/R209H*^ littermates following a 2.0 mg/kg dose of risperidone. Student’s unpaired *t-*test (B) At 0.5 mg/kg both WT and *Gnao1*^*+/R209H*^ exhibit a significant decrease in locomotion compared to vehicle controls. Student’s unpaired *t-*test. Wildtype mice also show a decrease in locomotion after 0.5 mg/kg risperidone treatment. (C) Comparison of 2.0 mg/kg and 0.5 mg/kg treatment in WT and *Gnao1*^*+/R209H*^ mice. Hyperactivity of *Gnao1*^*+/R209H*^ mice was observed during baseline testing and recovered following the 2-day risperidone washout.

## Discussion

We have developed a novel mouse model carrying an R209H mutation in the *Gnao1* gene that leads to a serious movement disorder (Ananth et al., 2016; Dhamija et al., 2016). While *GNAO1* mutations are rare in humans, the R209H mutation is the second most common found in patients with *GNAO1*-related disorders. The mice have a clear movement phenotype confirming the pathological nature of this mutation despite earlier work showing that it could normally support receptor-mediated regulation of cAMP in HEK cells (Feng et al., 2017). This model will provide a powerful tool to both understand neural mechanisms of the dysfunction of the R209H Gα_o_ protein as well as permitting preclinical drug repurposing studies.

The goal of personalized medicine is to define treatments for an individual patient. Knowing which gene is involved is beneficial but different mutant alleles, even in the same gene, can produce quite distinct effects. Here we report a second mouse model of *GNAO1*-related neurological abnormalities. The *Gnao1 R209H* mutant mice display motor abnormalities but no seizures which differs from our previous *Gnao1* G203R mouse model (Feng et al., 2019). This is consistent with the clinical pattern of *GNAO1* R209H patients, who present with neurodevelopmental delay with involuntary movements (NEDIM) without seizures (Ananth et al., 2016; Dhamija et al., 2016).

The specific movement abnormalities seen with the *Gnao1* R209H mutant mice were somewhat unexpected; our previous mouse model, *Gnao1*^*+/G203R*^, showed significant motor impairments in Rotarod and Digigait but had no changes in the open field test (Feng et al., 2019). In contrast, the *Gnao1* R209H mutant mice reported here have profound hyperactivity in the open field test and either no or only modest deficits in rotarod and digigait analysis. Patients with the G203R or R209H mutations both display movement disorder with choreathetosis and dystonia as the most common pattern (Feng et al., 2018; Saitsu et al., 2016). Interestingly, patients with the R209H mutation have somewhat more choreoathetosis compared to those with the G203R mutation, although they display similar frequencies of dystonia (Feng et al., 2018). Such differences in patient phenotype may relate to the differences in the animal models behavior. In looking at the behavior of the R209H model specifically, the striking hyperactivity of the male R209H mutant mice points to increased dopamine signaling in the striatum as a possible mechanism. Suppression of that hyperactivity with risperidone is consistent with that mechanism.

The two movement disorder-associated *GNAO1* mutations R209H and G203R also have different patterns in *in vitro* studies of cAMP regulation in HEK-293T cells (Feng et al., 2017). Interestingly, the Gα_o_ R209H mutant supports normal cAMP regulation and is expressed at normal levels both in HEK cells (Feng et al., 2017) and in the brain (this study). In contrast the G203R mutant is expressed at lower levels both in HEK cells (Feng et al., 2017) and in the mouse brain (Feng, H. Unpublished doctoral dissertation). Despite lower expression, it signals more strongly (GOF) in the HEK cell assay.

We also present two new pieces of information that provide insights into the pathogenicity of the R209H mutant Gα_o_ despite its normal function in the HEK cell cAMP assay. Using a fluorescent nucleotide exchange assay with pure R209H mutant Gα_o_, we demonstrate that the rate of binding of the GTP analog is ∼9x faster than that of WT. This is typical for many hyperactive G protein mutants since GDP release is the rate-limiting step for activation. Indeed, receptors enhance GDP release rates to activate the G protein (Iiri, Herzmark, Nakamoto, van Dop, & Bourne, 1994; Leyme, Marivin, Casler, Nguyen, & Garcia-Marcos, 2014; Toyama et al., 2017). Constitutive activation of Gα_o_ by the R209H mutant could lead to enhanced signaling in the indirect pathway, D2 dopamine receptor neurons in striatum. The striatum is a site of substantial Gα_o_ expression and enhanced Gα_o_ signaling could contribute to the hyperactivity phenotype seen in our studies. More detailed mechanistic studies of striatal neurons will be needed to further elucidate the downstream signaling mechanisms protein (e.g. ion channels, synaptic vesicle release, or neurite outgrowth) engaged by the mutant G_o_ in these mice and in patients.

Effective treatments are a key goal as patients with the R209H mutation experience repeated hospitalizations (Ananth et al., 2016; Marecos et al., 2018; Schirinzi et al., 2018). Deep brain stimulation in the internal globus pallidus has proven effective in *GNAO1* patients in attenuating MD (Honey et al., 2018; Koy et al., 2018; Kulkarni et al., 2016; Sanem et al., 2016). However, that invasive treatment is reserved for refractory patients. Risperidone is one of the oral treatments that has proven to be beneficial, specifically in a patient with the R209H mutation (Ananth et al., 2016). Risperidone is an atypical neuroleptic, antagonizing D_2_ and 5-HT receptors. Gα_o_ couples to myriad GPCRs including the dopamine D_2_ receptor which is involved in movement control (Neve, Seamans, & Trantham-Davidson, 2004). In our study, risperidone was able to significantly decrease the hyperlocomotion seen in our *Gnao1*^*+/R209H*^ mouse model. At both the 0.5 mg/kg and 2.0 mg/kg dose of risperidone, hyperactivity was attenuated in our R209H mouse model. However, this response was not selective for the *Gnao1*^*+/R209H*^ mutant mice as the WT mice also displayed a significant decrease in locomotion. This outcome suggests that risperidone treatment may be effective in repressing global movement, while not specifically targeting a Gnao1 R209H mechanism.

The new mouse model described here should provide a valuable tool for future mechanistic studies of *GNAO1* encephalopathies. The fact that the mouse has very distinct behavioral changes from our previous *Gnao1* mutant mouse models (R209H and G184S), indicates that it is very important to consider the mutant allele as well as the mutant gene in considering genetic disorders and personalized therapies.

## Abbreviations

AC: Adenylyl Cyclase
cAMP: Cyclic adenosine monophosphate
EIEE17: Early infantile epileptic encephalopathy 17
GOF: Gain-of-function
GPCR: G-protein coupled receptor
GTPγS: 5’-Guanosine-diphosphate-monothiophosphate
LOF: Loss-of-function
NEDIM: Neurodevelopmental disorder with involuntary movements
RNP: Ribonucleoprotein
tracrRNA: Trans-activating crRNA
crRNA: CRISPR RNA
ssODN: Single-stranded oligodeoxynucleotide
DSB: Double strand DNA break
PAM: Protospacer-adjacent motif

## Acknowledgements

The authors thank Dr. Michelle Mazei-Robison for her advice on organizing the behavioral battery experiments on our mutant mouse models.

## Author Contributions

Data Curation: Casandra L. Larrivee, Huijie Feng, Josiah A. Quinn, Vincent S. Shaw, Elena Y.

Demireva, Huirong Xie, Jeff R. Leipprandt

Formal Analysis: Casandra L. Larrivee, Huijie Feng, Josiah A. Quinn, and Vincent S. Shaw.

Methodology: Casandra L. Larrivee, Huijie Feng, Elena Y. Demireva, Josiah A. Quinn, Vincent S.

Shaw, Richard R. Neubig.

Resources: Jeff R. Leipprandt.

Supervision: Casandra L. Larrivee, Huijie Feng, Richard R. Neubig.

Visualization: Huijie Feng.

Writing – original draft: Casandra L. Larrivee

Writing – review & editing: Casandra L. Larrivee, Huijie Feng, Richard R. Neubig

