## Supplemental File for "Mice with *GNAO1* R209H Movement Disorder Variant Display Hyperlocomotion Alleviated by Risperidone"

**Supplemental Materials**

**Table S1. *GNAO1* R209H Patient Classification**

| **Patient No.** | **Sex** | **Amino Acid Change** | **Age of Onset** | **Presence of Epilepsy** | **Movement Disorder** | **Treatment** | **Motor Developmental Delay(MDD)/Intellectual Delay(ID)** | **Reference** |
| --- | --- | --- | --- | --- | --- | --- | --- | --- |
| 1 | M | R209H | 17 mo | - | Chorea | DBS | MDD | Kulkarni et. al (2016) |
| 2 | M | R209H | 2 y | - | Chorea | DBS | MDD | Kulkarni et. al (2016) |
| 3 | M | R209H | 3 y | - | Chorea | Risperidone, BZD | MDD/ID | Anath et. al (2016) |
| 4 | M | R209H | 1 y |  | Chorea | NA | MDD/ID | Menke et al (2016) |
| 5 | M | R209H | 10 mo | - | Chorea Dystonia | TBZ, THP | MDD/NA | Dhamija et al (2016) |
| 6 | M | R209H | 15 mo | - | Chorea, Dystonia | DBS | MDD/ID | Marecos et al (2018) |
| 7 | F | R209H | 6 mo | - | Dystonia | NA | MDD/MID | Kelly et al (2018 |
| - | F | R209C | NA | NA | Chorea | NA | MDD/ID | Saitsu et al (2016) |
| - | F | R209G | 3 y | - | Chorea | None | MDD/ID | Anath et al (2016) |
| - | M | R209L | 2 y | - | Chorea | NA | MDD/ID | Menke et al (2016) |

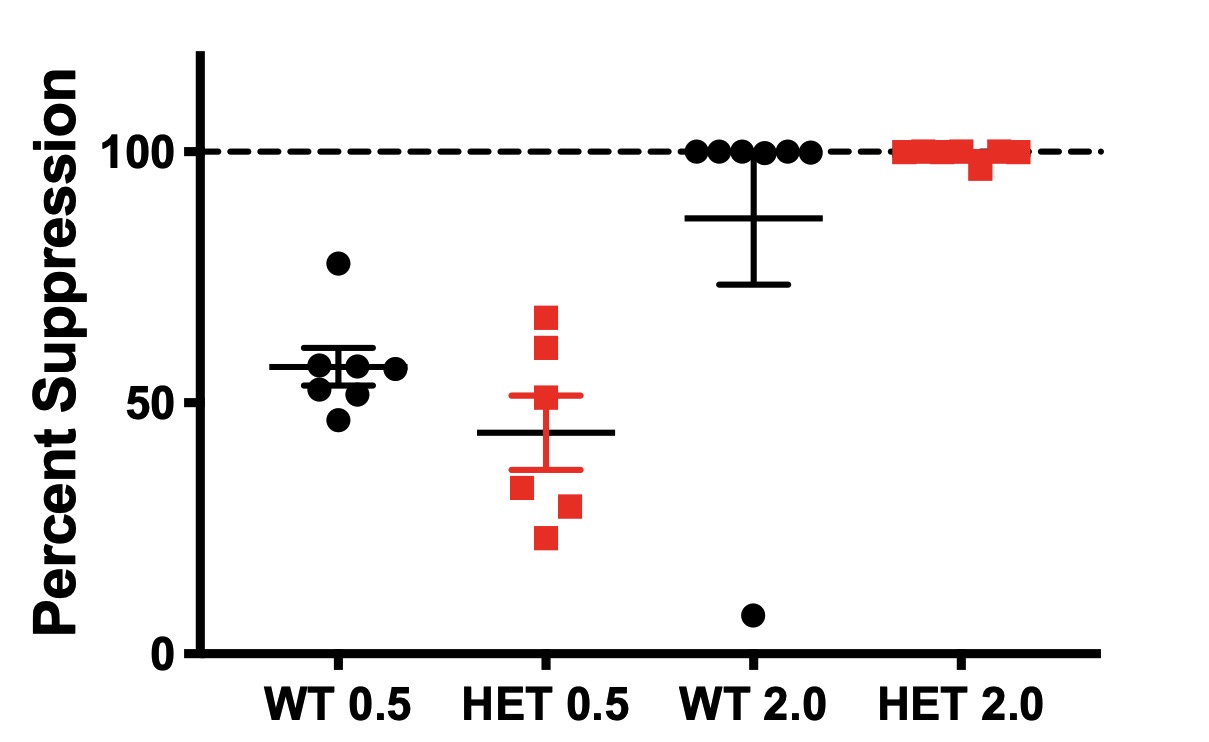

**Figure S1. WT and *Gnao1^+/R209H^* mice show no difference in percent suppression of locomotion after risperidone treatment** *Gnao1^+/R209H^* mice show similar sensitivity to risperidone treatment at 2.0 mg/kg or 0.5 mg/kg compared to WT treated mice, unpaired Student’s *t*-test.

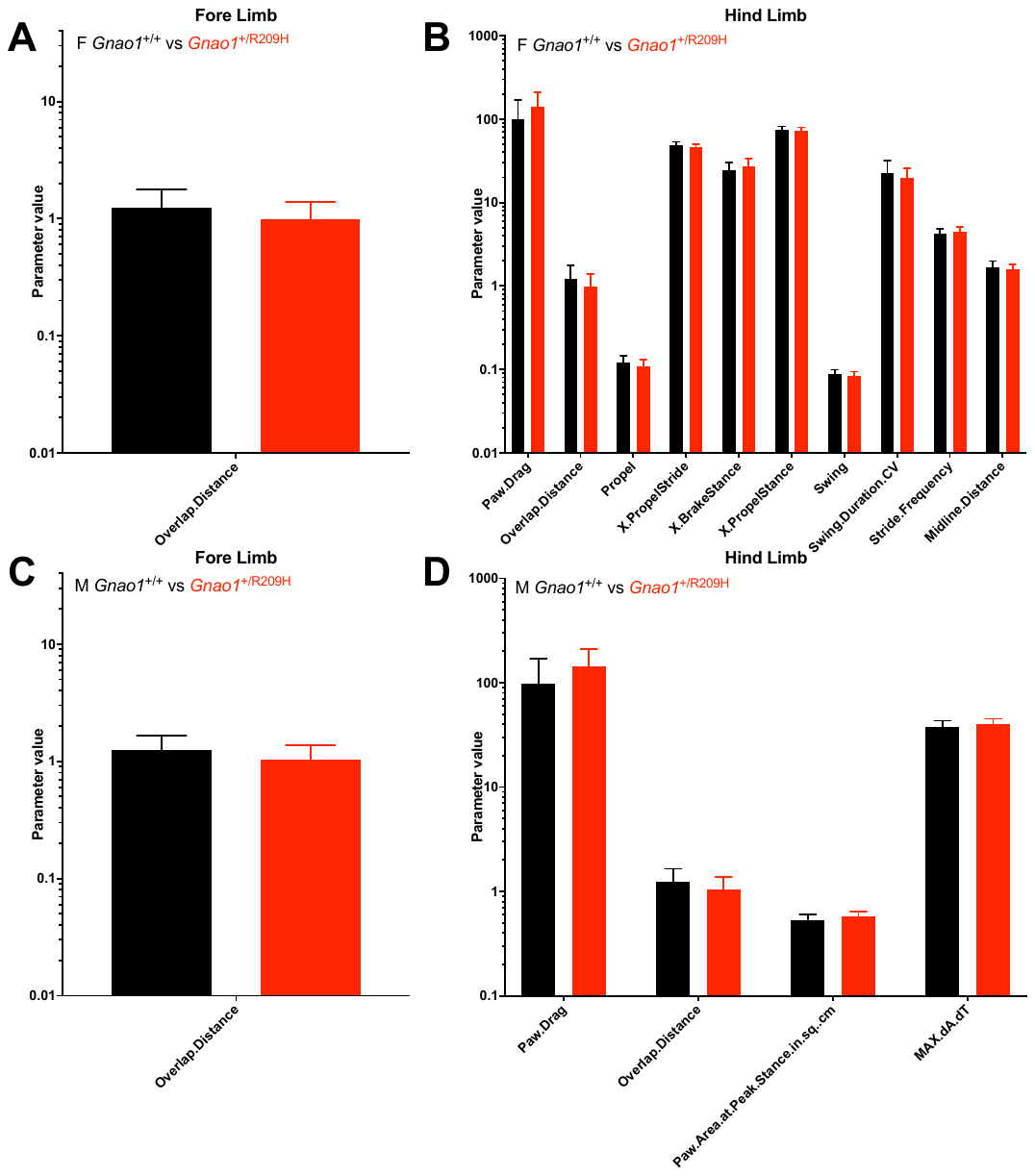

**Figure S2. False discovery rate (FDR) calculation probed of significantly different parameters from the DigiGait data in *Gnao1*^+/R209H^ mice.** All parameters that showed significance are plotted here. (A&B) Female *Gnao1*^+/R209H^ and their littermate controls showed 9 parameters with significance detected by the FDR analysis. (C&D) Male *Gnao1*^+/R209H^ and their littermates controls exhibited fewer parameters with significance comparing to female detected by the FDR analysis in fore and hind limb data combined. FDR is calculated by a two-stage step-up method of Benjamini, Krieger and Yekutiel. Significant values are defined as q < 0.01.

**Table S2. Gait analysis parameters of female *Gnao1* R209H mutant mice**

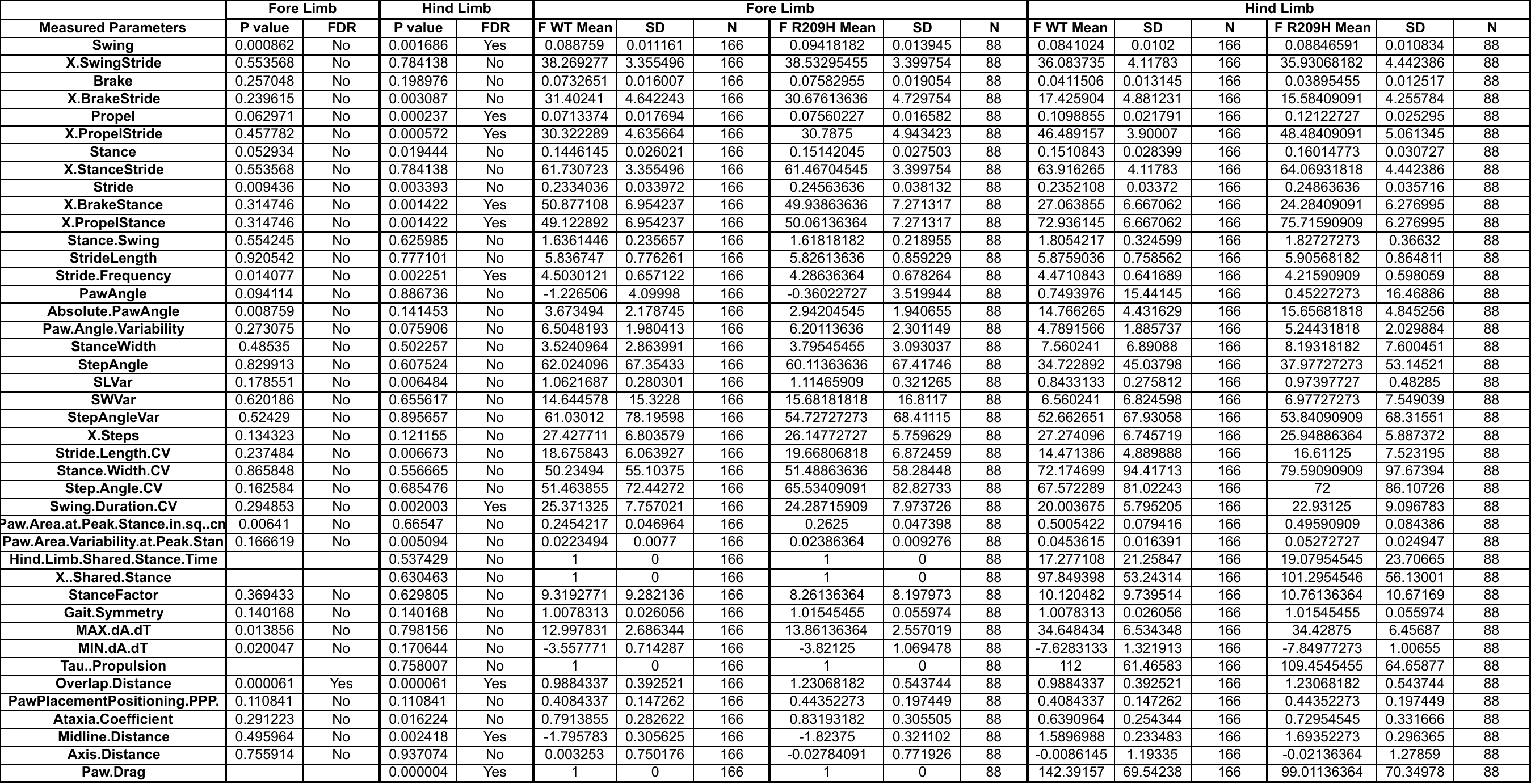

**Table S3 Gait analysis parameters of male *Gnao1* R209H mutant mice**

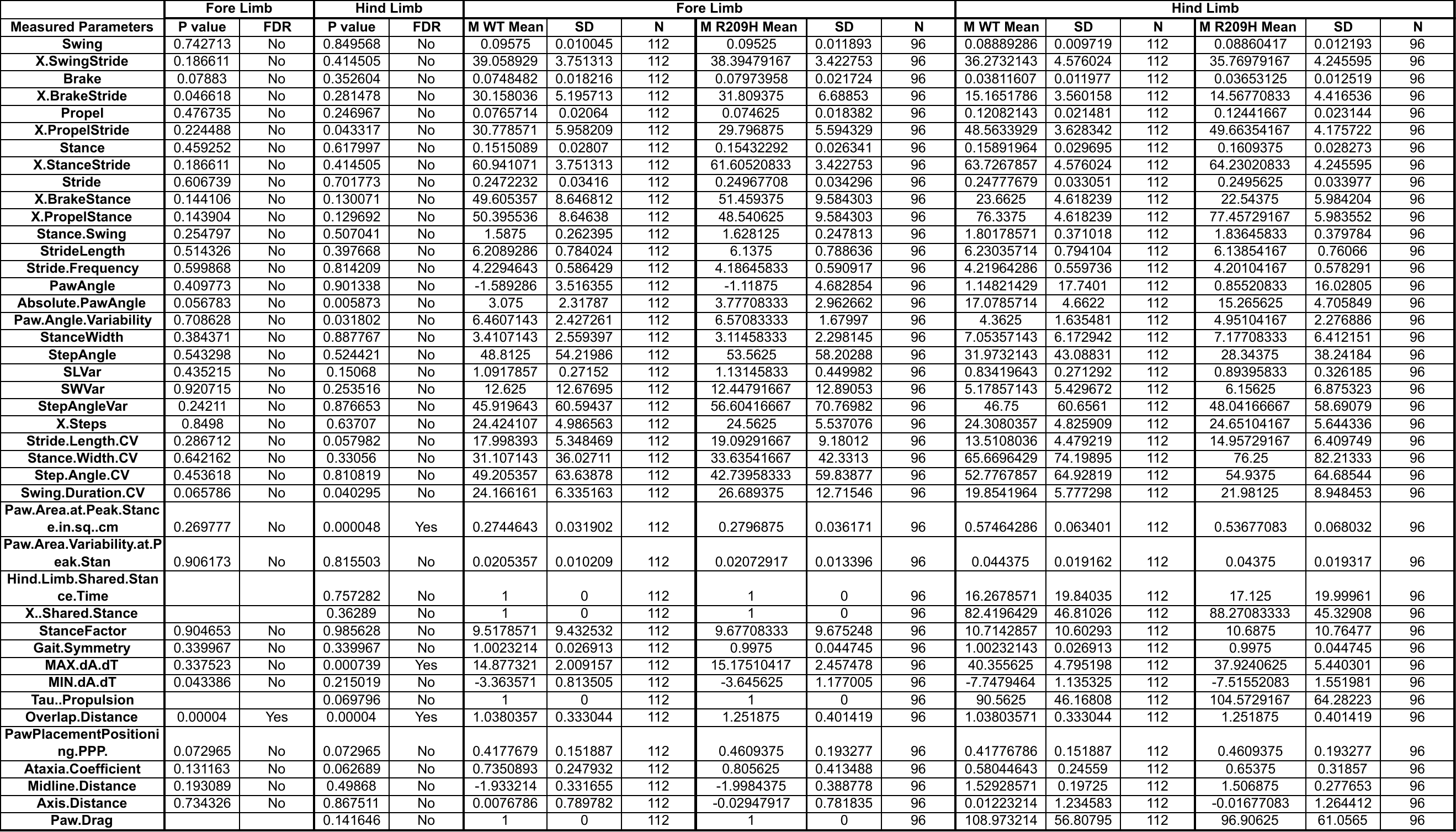
